# Structure-based discovery of inhibitors of Mac1 domain of nonstructural protein-3 of SARS-CoV-2 by machine learning-augmented screening of chemical space

**DOI:** 10.1101/2025.09.05.674529

**Authors:** Fuqiang Ban, Rahul Ravichandran, Galen J. Correy, Oleksandra Herasymenko, Madhushika Silva, Suzanne Ackloo, Albina Bolotokova, Irene Chau, Elisa Gibson, Rachel Harding, Ashley Hutchinson, Peter Loppnau, James S. Fraser, Matthieu Schapira, Artem Cherkasov, Francesco Gentile

**Author notes:** (AC), (FG). Equal contribution (Fuqiang Ban, Rahul Ravichandran).

## Abstract

Significant efforts have been recently dedicated to the discovery of small molecule inhibitors against the Macrodomain 1 (Mac1) of nonstructural protein 3 (NSP3) as potential antivirals for SARS-CoV-2. Thus, Mac1 has also been selected as the target for the Critical Assessment of Hit-finding Experiments (CACHE) challenge #3. As contestants in that challenge, we developed a computational strategy that ranked on the top among all 23 participants in the competition and resulted in the discovery of a novel chemical series of non-charged Mac1 inhibitors. Those have been identified through the combination of machine learning-accelerated virtual screening of Enamine REAL Diversity Subset of approximately 25 million compounds and consequent hit expansion into the entire Enamine REAL Space library. In particular, the initially identified hit compound CACHE3-HI_1706_56 (K_D_ = 20 µM) was explored by probing 17 close analogues from a library of 44 billion molecules from the Enamine REAL. All those analogues effectively displaced the Mac1-binding ADP-ribose peptide, and 12 were confirmed to engage with Mac1 by the Surface Plasmon Resonance experiments, revealing a new chemical series of compounds for hit-to-lead optimization.

The structure of the CACHE3-HI_1706_56-Mac1 complex was further determined at high resolution with crystallography, confirming initial computational predictions. Our results illustrate the effectiveness of ML-accelerated docking to rapidly identify novel chemical series and provide a strong foundation for the development of SARS-CoV-2 NSP3 Mac1 inhibitors.

## Introduction

ADP-ribosylation is a NAD+-dependent modification of proteins or nucleic acids, broadly categorized as Mono(ADP-ribosyl)ation (MARylation) or poly(ADP-ribosyl)ation (PARylation). The process of MARylation plays a complex role in antiviral response through modifying host and viral proteins. To counteract antiviral ADP-ribosylation by host Poly(ADP-ribose) polymerases (PARPs), SARS-CoV-2 and other coronaviruses encode a conserved macrodomain within the nonstructural protein 3 (NSP3) that hydrolyzes ADPribose modifications, thereby reversing the effect of ADP ribosylation^1^, which enhances interferon signalling^2^. In SARS-CoV-2, the Macrodomain 1 (Mac1) of NSP3 plays a key role in the immune evasion capabilities responsible for the coronavirus disease 2019 (COVID-19) pandemic^3^. In particular, Mac1 displays mono(ADP-ribosyl) hydrolase activity to block interferon response and acts against the host defense mechanism, which is essential and critical for coronavirus pathogenesis and lethality^1, 4^. An active site mutation inactivating Mac1 has been shown to render SARS-CoV-1 and SARS-CoV-2 nonlethal in a lethal mouse model of viral infection, and virus bearing the mutation do not replicate as efficiently in interferon-stimulated cells as wild type virus.^5^ Therefore, Mac1 has emerged as a potential antiviral drug target with promising therapeutic value.

Following the onset of COVID-19, many inhibitors of Mac1 have been reported.^6^ In particular, Gahbauer et al. identified 160 Mac1 ligands representing 119 different scaffolds and determined 152 complex crystal structures through iterations of fragment screening, structure-based design, protein crystallography, and binding evaluation, identifying several molecules with low- to sub-micromolar potencies.^7^

Two other approaches of fragment-linking^8^ and virtual screening of ultra-large chemical libraries^9^ were also employed for the discovery and optimization of potent Mac1 ligands. The fragment-linking method leveraged the crystallographic structures of 234 fragment-Mac1 complexes that were previously determined^8^ by merging pairs of fragments binding to different subpockets to generate full-size molecules. An automated fragment merging and linking strategy, paired with SAR-by-ultra-large catalogue optimization led to the discovery of compound Z8539_0072 (Figure 1A) with an IC_50_ of 0.4 μM which complex with Mac1 was further determined by crystallography (pdb 5SQW). In parallel, virtual screening of ultra-large and focused chemical libraries of lead-like molecules were conducted against the closed and everted state of Mac1 (depending on the conformation of the Ala129 to Pro136 loop), from which both neutral and anionic inhibitor scaffolds were identified and optimized. Although anionic series led to potent molecules, poor cell permeability represented a challenge. For example, the Z8539 scaffold had low cell permeability (11 nm/s) in MDCK cells that limits its potential antiviral activity.^7^ Thus, non-charged inhibitors were also designed by installing neutral hydrogen bond acceptors in the oxyanion hole pocket, resulting in the most active neutral inhibitor LRH-0003 (pdb 5SRY, IC_50_ 1.7 µM) shown in Figure 1B, with high permeability values.

**Figure 1.**
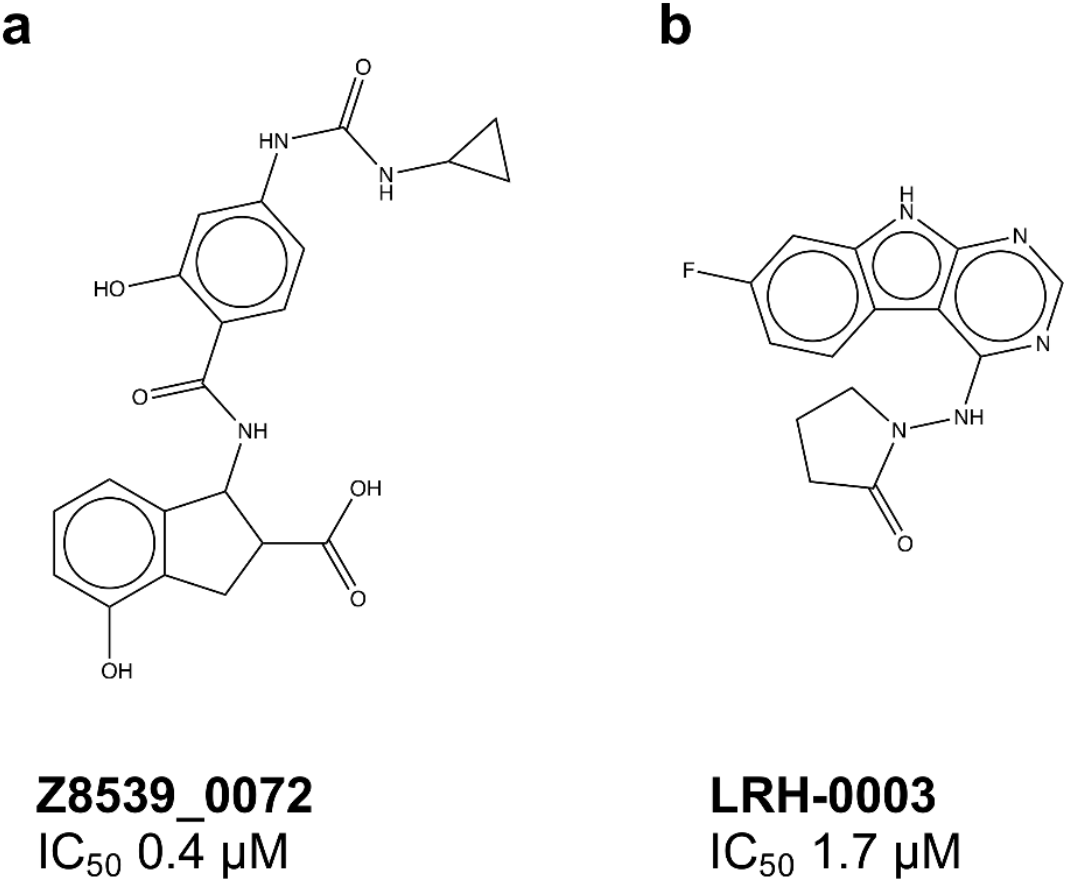
Chemical structures of SARS-CoV-2 NSP3 Mac1 inhibitors. a) best anionic inhibitor, Z8539_0072 b) Best neutral inhibitor, LHR-0003^7^.

The Critical Assessment of Computational Hit-finding Experiments (CACHE) challenge #3 is part of a series of prospective benchmarking exercises to evaluate progress in the field of computational hit finding that are open to both academic and industrial groups^10^. In CACHE3, the main goal was to identify novel Mac1 inhibitor scaffolds able to compete with the ADP ribose substrate without bearing negatively charged moieties. The challenge comprised two competitive rounds. In the first round, computational chemistry experts were invited to select ∼ 100 compounds from commercial libraries as potential binders of SARS-CoV-2 NSP3 Mac1. The compounds were then purchased and experimentally tested for their capacity to competitively inhibit ADP ribose binding. All positive hits were then advanced to Round 2, a hit expansion round where participants selected ∼ 50 follow-up molecules for chemical series confirmation. Based on both rounds of CACHE-3, an independent committee composed of industry experts assessed the validity of the biophysical activity data of each series, the drug-likeness of the validated hits, and their suitability as starting points for hit-to-lead optimization, providing a final rank associated with each participating workflow.

In this study, we present our CACHE-3 winning workflow that enabled to discover novel Mac1 inhibitor chemotypes within ultra-large chemical databases. Specifically, we utilized Deep Docking-accelerated virtual screening^11-12^ coupled with expert selection^13^ and consensus strategies^14^ to identify a novel scaffold within a diversified subset of Enamine REAdily accessibLe (REAL) database. Computational analogue search in the vast REAL Space led to the efficient identification of a neutral, novel chemical series of inhibitors engaging Mac1 with low µM affinities. X-ray crystallography confirmed the predicted binding mode of the best hit compound, providing the basis for future efforts in developing cell-permeable Mac1 inhibitors.

## Materials and Methods

### Protein structure preparation

Pdb 5SRY, corresponding to the crystallographic LRH-0003-Mac1 complex determined at a resolution of 1.05 Å^7^, was used as target structure for virtual screening. The structure was obtained from the Protein Data Bank (PDB)^15^ and prepared with the standard procedure of Protein Preparation Wizard tool from Maestro GUI of Schrödinger suite of programs^16^. Water molecules and heteroatoms were removed in the final step.

### Small molecule library preparation

The SMILES of 43,759,934 compounds in the Enamine REAL Diversity Subset^17^ were downloaded, and chirality was assigned using OpenEye’s flipper tool^18^. Structures with more than two chiral centers were removed. OpenEye’s tautomers tool^19^ was used to calculate one dominant ionization state per molecule. Prior to docking, one 3D low-energy conformation per small molecule was generated with OpenEye’s omega tool in *classic* mode^18^.

### Docking

Docking grids were centered on LRH-0003 and individually set up in Maestro and ICM^20^ GUIs. Docking was performed using Glide Standard Precision (SP)^21^, and ICM^20^, using the default settings.

### ML-accelerated virtual screening

The Deep Docking protocol^12^ was used for the virtual screen, with a recall of 0.9 for the positive class. Molecules were represented as Morgan fingerprints with radius of 2 and 1,024 bits^22^ computed with RDKit^23^. Five iterations were run using Glide SP as docking engine with an initial training/validation/test size of 50,000 molecules each, and 50,000 molecules added to the training at each subsequent active learning iteration. The percentage of top scoring molecules classified as positive was linearly decreased from 1 to 0.1% across the five iterations.

### Hit selection

A first set of potential hits with Glide SP score equal or less than −8 kcal/mol was redocked with ICM, and subsequently filtered using a two program consensus filter^24-25^ with a root-mean square deviation (RMSD) cutoff of 2 Å, followed by a consensus scoring protocol based on three scores: Glide SP score, ICM score, and Molecular Operating Environment (MOE) pKi score computed with the *scoring*.*svl* script. A vote of 1 in each category was assigned if the molecule belonged to the top 10% set based on its score, and molecules were ranked based on the combined score. In parallel, top scoring molecules from the Glide SP and ICM raw ranks were visually inspected, and a second set of potential hits was identified.

### Chemical series expansion

BiosolveIT InfiniSee^26^ was used to identify analogues of the hit confirmed in the first step of the CACHE challenge, within the Enamine REAL Space (44 billion compounds)^27^. Retrieved molecules were prepared as before and then docked to Mac1 with Glide SP. Visual inspection was used to shortlist 75 promising candidates for quotation.

### Protein expression and purification

DNA fragments encoding SARS-CoV-2 NSP3 Macro 1 domain (residues 205–380), NSP3 Macro 1 domain (206–374), and human PARP14 (residues 994–1196) were subcloned into expression vectors with N-terminal His tags. Specifically, NSP3 (205– 380) was cloned into pDEST17, while NSP3 (206–374) and PARP14 (994–1196) were inserted into pNicBio3 and pNIC-Bio2, respectively, with C-terminal AviTags for biotinylation.

Protein constructs were expressed in *E. coli* BL21 (DE3) or BL21 (DE3)-BirA (for AviTag constructs) in Terrific Broth supplemented with antibiotics. Cultures were induced at OD_600_ 0.8–1.5 with 0.5 mM IPTG and incubated overnight at 18 °C. For AviTag constructs, D-Biotin (10–100 µM) was added to the media.

Cells were harvested and lysed in Tris-HCl buffer (pH 7.5) containing 500 mM NaCl, imidazole, glycerol, and a protease inhibitor cocktail (Aprotinin, Leupeptin, Pepstatin A, E-64). Chemical lysis was performed using CHAPS, TCEP, PMSF/Benzamidine, and Benzonase, followed by sonication (5–10 min, Sonicator 3000, Misoni). The lysates were clarified by centrifugation (36,000 ×g, 60 min, 4 °C).

Purification was performed as follows: NSP3 (205–380), Ni-NTA affinity chromatography eluted with imidazole, followed by gel filtration (HiLoad Superdex75 26/600, ÄKTA Pure) in 50 mM Tris (pH 7.5), 250 mM NaCl, 0.5 mM TCEP, and 5% glycerol; NSP3 (206–374), Ni-NTA affinity chromatography with a pre biotin wash and eluted with imidazole, followed by gel filtration under the same conditions; and PARP14 (994–1196) Ni-NTA affinity chromatography with a pre biotin wash and eluted with imidazole, followed by dialysis in 20mM Tris, pH 8, 500mM NaCl, 1mM TCEP.

All three proteins were purified to 95% purity, assessed by SDS-PAGE, pooled, concentrated, snap-frozen, and stored at −80 °C. Protein identity was confirmed by LC-MS.

### Homogeneous Time-Resolved Fluorescence (HTRF)

Binding affinity of the tested compounds to SARS2 NSP3 Mac1 (205-380) protein was assessed by the displacement of an ADP-ribose conjugated biotin peptide from His6-tagged protein using HTRF-based assay. Compounds were dispensed into ProxiPlate-384 Plus (PerkinElmer) assay plates using an Echo 650 acoustic liquid handler (Beckman Coulter). Binding assay was conducted in a final volume of 20 μl with 12.5 nM NSP3 Mac1 protein, 200 nM peptide ARTK(Bio)QTARK(Aoa-RADP)S (Cambridge Peptides), 0.031 nM Terbium-cryptate anti-His Mab (HTRF donor, PerkinElmer) and Streptavidin-XL665 (HTRF acceptor, PerkinElmer) in assay buffer (25 mM HEPES pH 7.0, 20 mM NaCl, 0.05% bovine serum albumin and 0.05% Tween-20). Assay reagents were dispensed into plates using a Multidrop Combi (ThermoFisher Scientific). Macrodomain protein and peptide were first dispensed and incubated with the tested compounds for 30 min at room temperature, followed by addition of the HTRF reagents and incubation at room temperature for 1 h. Fluorescence was measured using a Synergy H1 microplate reader (Biotek) with the HTRF filter set (A = excitation 330/80 nm, emission of 620/10 nm, and B = excitation of 330/80 nm and emission of 665/8 nm). Obtained HTRF ratio values were used to estimate the percentage of ADP-r peptide binding inhibition/displacement using a positive control (4% DMSO solution in presence of protein and peptide) and negative control (4% DMSO solution in presence of peptide) values from each screening plate.

### Surface Plasmon Resonance (SPR)

Orthogonal binding confirmation was assessed by SPR using streptavidin-conjugated (SA) chip. The assay was conducted using a Biacore™ 8K (Cytiva) instrument at 20°C. Biotinylated SARS2 NSP3 (206-374) protein was immobilized onto the active flow cell of the SA chip following the manufacturer’s protocol reaching approximately 2800-3200 response units (RU). For the counter screen phase half of the channels were charged with unrelated negative control protein PARP14 (994-1196) reaching approximately 2800-3000 RU. All were kept empty, and their response was subtracted for each channel respectfully. Compounds were initially dissolved in 100% DMSO to create 10 mM stock solutions, which were subsequently serially diluted (factor: 0.5) to obtain six concentrations points in 100% DMSO. For the SPR run, these serially titrated compound stocks were diluted 1:25 in HBS-EP+ buffer (10mM Hepes pH7.4, 150mM NaCl, 3mM EDTA, 0.05% (v/v) Tween20) supplemented with 0.5 mM reducing agent (TCEP) to achieve a final concentration of 4% DMSO. Binding experiments used multicycle kinetics with a contact time of 60 seconds and a dissociation time of 120 seconds at a flow rate of 40 µL/min at 20°C. The dissociation constant (K_D_) values were determined using steady-state affinity 1:1 binding with Biacore™ Insight Evaluation software (Cytiva).

### Dynamic Light Scattering

The solubility of compounds was estimated by DLS that directly measures compound aggregates and laser power in solution. Compounds were prepared at 2.5 mM and 1.25 mM directly from DMSO stocks, then diluted 25x into filtered 10mM Hepes pH7.4, 150mM NaCl, 3mM EDTA, 0.5 mM TCEP (4% DMSO final). The resulting samples were then distributed into 384-well plates (Corning, Cat# 3540), with 20 μL in each well. The sample plate was centrifuged at 3500 rpm for 5 min before loading into DynaPro DLS Plate Reader III (Wyatt Technology).

### Crystallography

The construct of Mac1 that crystallizes in P43 was expressed, purified and crystallized as described previously^8^. Ligands prepared in DMSO at 50 mM were soaked into crystals achieving a nominal concentration of 5-10 mM using acoustic dispensing^28^. After 2-4 hours at room temperature, crystals were vitrified in liquid nitrogen with assistance from a Crystal Shifter^29^. X-ray diffraction data were collected at beamline 8.3.1 (supplementary X-ray statistics). Data were indexed, integrated and scaled using XDS^30^ and merged with Aimless^31^. After initial rigid body refinement with phenix.refine^32^, coordinates were refined using Refmac5^33^ as described previously^34^. Ligands were identified and modeled using PanDDA^35^ run in CCP4 version 7.0^33^. Ligand coordinates and restraints for refinement were generated with LigPrep (version 2022-1, Schrodinger) and phenix.elbow^36^ or Grade2 (version 1.7.0, Global Phasing Limited). Because of the low occupancy of the ligand, coordinates were refined using phenix.refine as a multi-state model with the apo state assigned alternative location identified (altloc) A and the ligand-bound state assigned altloc B as described previously^37^.

## Results and Discussion

### ML-accelerated virtual screening against Mac1

At the beginning of the CACHE3 challenge, there were more than 160 Mac1 crystal structures available in the PDB database. After analyzing the corresponding structures and considering the restriction of predicting only neutral inhibitors, the crystal structure of the most potent neutral inhibitor LRH-0003 (IC_50_ = 1.7 µM) in complex with Mac1 was selected for our docking campaign. LRH-0003 was specifically designed to interact with the oxyanion hole (NH groups of Phe156 and Asp157) of Mac1 through a neutral 1-aminopyrrolidin-2-one group^7^. Its potency is comparable to one of the most active anionic compounds (Z8539) as well as of an analogue carrying a carboxylic group (LRH-0021)^7^suggesting that the development of neutral Mac1 inhibitor with high potency is feasible. The binding mode of LRH-0003 is illustrated in Figure 2a, highlighting one hydrogen bond with the side chain of Asp22, two hydrogen bonding interactions to the backbone amine of Ile23 and Phe156 as well as a weak one with backbone amine of Asp157, and an important hydrophobic interaction with Phe156. Structurally, LRH-0003 engages Mac1 similarly to charged inhibitors, with the pyrrolidinone occupying the “oxyanion hole” generated by the backbone amines of Phe156 and Asp157.

**Figure 2.**
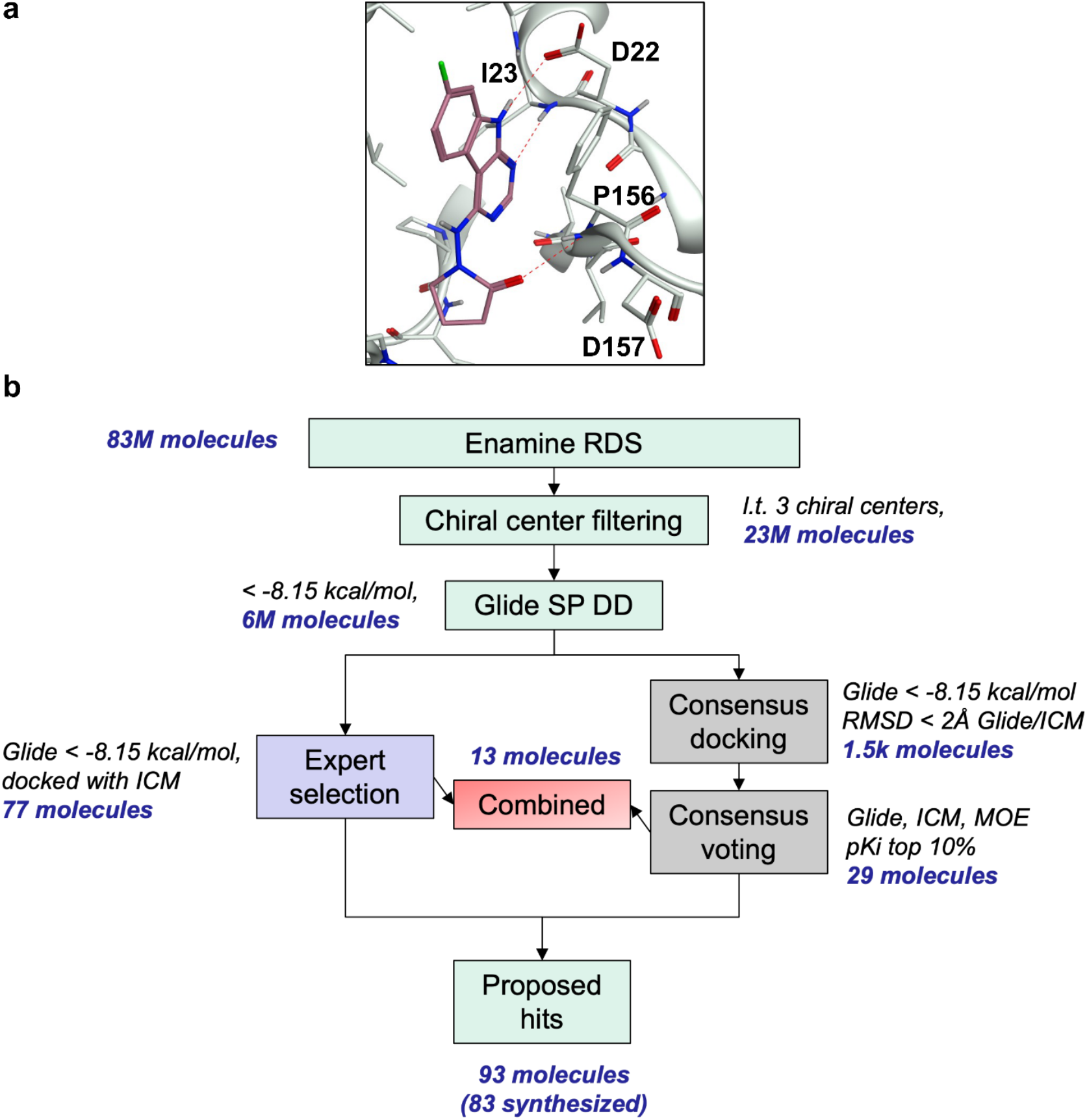
Binding site and virtual screening schematics. a) X-ray binding pose of LHR-0003 bound to SARS-CoV-2 NSP3 Mac1 (PDB 5SRY), the complex that was chosen as the target structure for virtual screening b) Multi-step structure-based virtual screening workflow to investigate the Enamine REAL Diversity Subset against Mac1.

Next, we employed the machine learning (ML)-accelerated virtual screening workflow to identify potential candidates for experimental validation (Figure 2b). First, we filtered out molecules with more than two chiral centers from the Enamine REAL Diversity Subset and expanded stereochemically the remaining entries, resulting in 23,139,109 molecules in total. Within the Deep Docking active learning framework, we iteratively docked 350,000 molecules (∼1.5% of the entire library) into the Mac1 binding site, divided in 50,000 for validation and test set and 50,000 for the training pool in each iteration, for five iterations. Through the process, the library was iteratively pruned of low-scoring molecules, resulting in 6,571,192 final molecules that included 90% of the top 0.1% scoring hits, that is, molecules with a Glide SP score equal or less than −8.15 kcal/mol. Notably, the reduction power of Deep Docking was significantly lower than the one that is usually observed for larger libraries. This is consistent with the high recall value that we used, the limited size of the library, and the diverse nature of the molecules within it, since structural congestion of full make-on-demand libraries is less relevant to diversity subsets. Nevertheless, the library was reduced by 3.5-fold while retaining the vast majority of top scoring molecules, by physically docking only 1% of the molecules.

The 6.5 million molecules were then docked to Mac1 with Glide SP. 26,057 molecules with a score equal to or less than −8 kcal/mol were retained and docked with ICM to the same site. RMSD-based consensus resulted in 1,553 molecules with Glide and ICM poses differing of no more than 2 Å. 29 molecules were prioritized for procuring and testing based on consensus scoring. In parallel, visual inspection of Glide and ICM docked sets was conducted, resulting in 77 prioritized molecules. 13 molecules were shared between the two sets, resulting in a final list of potential hits of 93 compounds. 83 compounds (89%) were obtained from Enamine, including 26 consensus- and 69 expert-selected molecules (12 molecules common to both sets).

### Discovery and characterization of a novel Mac1 inhibitor scaffold

Out of the 83 tested compounds, 11 inhibited the binding of the ADP-ribose peptide to Mac1 in the HTRF assay 100 µM concentration. Eight of them displayed dose-response inhibitory activity and passed the DLS solubility evaluation. Finally, only one molecule was selected as a hit (CACHE3-HI_1706_56, 4-[(2-bromocyclohex-2-en-1-yl)amino]-7H-pyrrolo[2,3-d]pyrimidine-5-carboxamide, Figure 3a), as it was confirmed to bind NSP3 Mac1 with SPR with a K_D_ of 20 µM (Figure 3b), without binding to PARP14a and without showing aggregation up to 100 µM.

**Figure 3.**
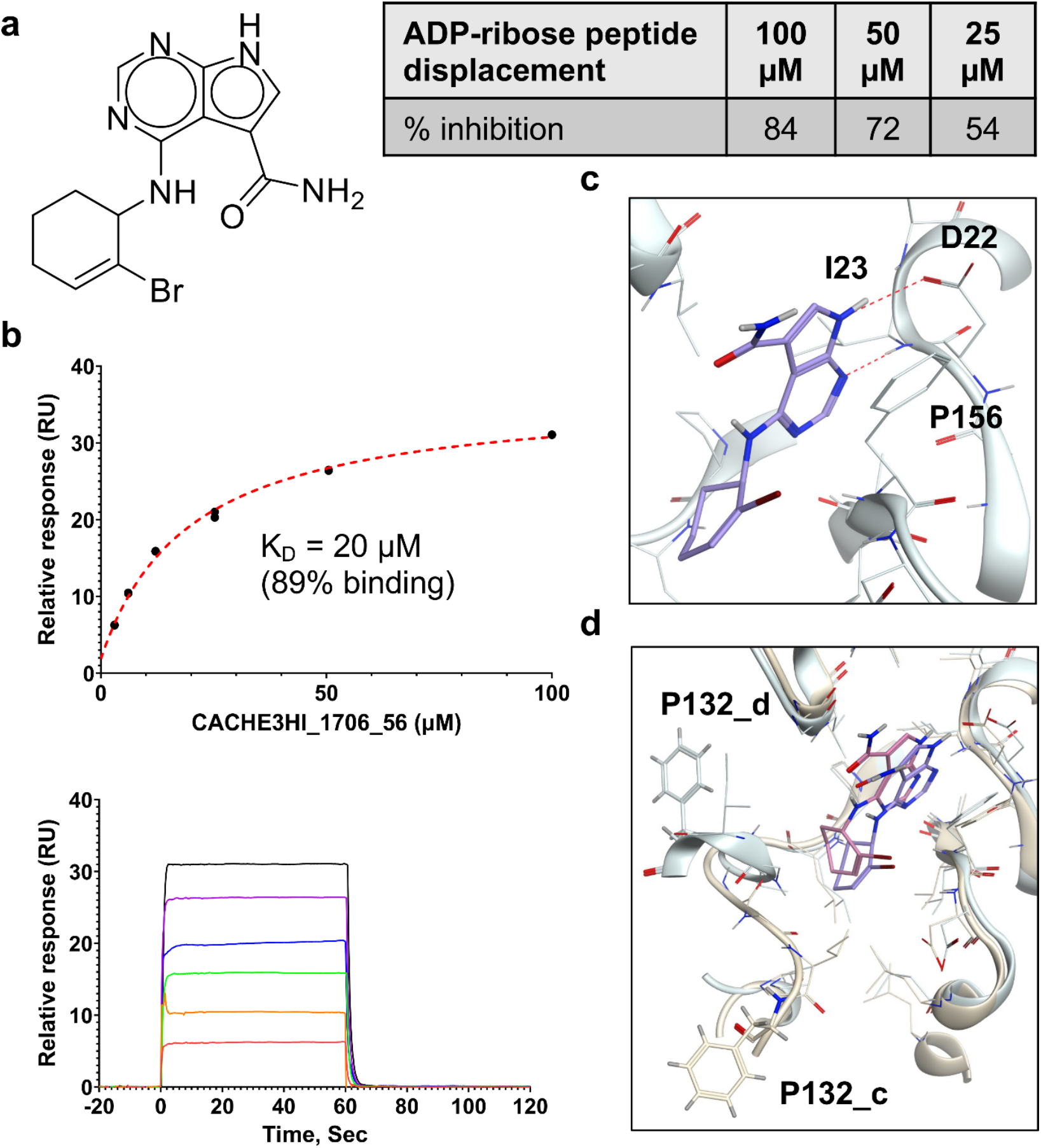
Experimental confirmation of the Round 1 hit. a) Chemical structure and HTRF results demonstrating inhibition of binding of the ADP-ribose peptide to Mac1 by CACHE3-HI_1706_56 b) Biophysical confirmation of Mac1-CACH3-HI_1706_56 binding with SPR c) Docking-predicted binding pose of CACHE3-HI_1706_56, with hydrogen bonds represented in red dashed lines d) a) Comparison between docked (ligand in blue, Mac1 in light green) and crystallographic (ligand in pink, Mac1 in yellow) Mac1-CACHE3-HI_1706_56 complexes; ligand binding induced a conformational change from the closed conformation of Phe132 (P132_d) used for docking and observed also in the apo-structure of Mac1 to the solvent-exposed one revealed by crystallography (P132_c). Only one of the two alternative conformations of CACHE3-HI_1706_56 is showed.

The binding pose of the hit predicted by Glide is illustrated in Figure 3c, highlighting hydrogen bonding between the 7H-pyrrolo[2,3-d]pyrimidine and the side chain of Asp22 and the backbone amine of Ile23. Interestingly, CACHE3-HI_1706_56 was selected as a potential hit in both the consensus and visual selection sets. Overall, the final hit rate of our virtual screening workflow was rather low (total 1/83 - 1.2% hit rate, 1/26 - 3.84% hit rate for consensus set, 1/69 - 1.45% hit rate for visually selected set, 1/12 - 8.33% hit rate for common set), highlighting the low computational tractability for the Mac1 site that was previously reported and echoed by the performances observed across all CACHE3 participants (31/1,739 confirmed hits, 1.8% hit rate).

The crystal structure of CACHE3_HI_1706_56 bound to Mac1 was solved with a resolution of 1.02 Å and two alternative, similar conformations, closely matching the computationally predicted pose. Interestingly, while the Mac1 conformation used in the virtual screen represented the closed state with respect to the conformation of the critical Phe132 residue, the crystal structure of the complex revealed that CACHE3_HI_1706_56 stabilizes instead the everted conformation with a noticeable displacement and solvent exposure of Phe132 (Figure 4a). This is consistent with previous observations that Mac1 hits can stabilize the everted conformation even if they were discovered by screening against the closed state, and vice versa^7^. The two conformations of CACHE3-HI_1706_56 both displayed hydrogen bonding interactions with the sidechain of Asp22 and backbone amine of Ile23, and an intramolecular hydrogen bonding between the carbonyl and NH linker of bromo-cyclohexene, as well as significant hydrophobic interactions with Ile131, Val149, and Phe156. In the oxyanion hole, the carbonyl of LRH-0003 was engaged in a hydrogen bond with backbone NH of Asp157, while CACHE3-HI_1706_56 did not display any hydrogen bonding interaction there. However, the bromine group was well tolerated. Finally, the amide group, while not displaying direct hydrogen bonding in the crystal structure, features the NH2 in proximity of the backbone carbonyl of Gly48.

**Figure 4.**
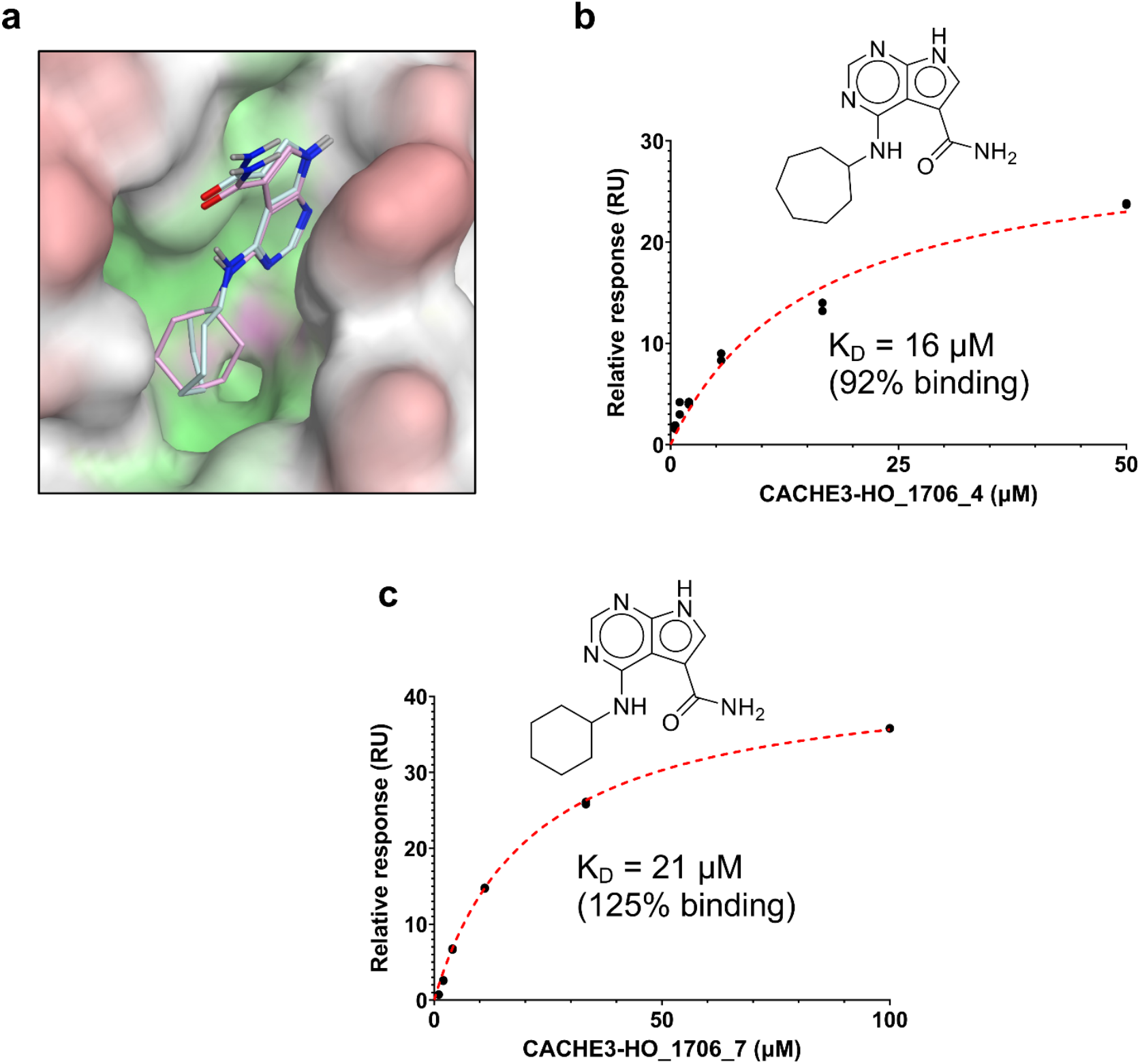
Binding pose and biophysical characterization of best hit analogues. a) Docking poses of CACHE3-HO _1706_4 (cyan) and CACHE3-HO _1706_7 (pink) to Mac1 pocket, represented as a molecular surface (green: hydrophobic regions, purple: polar regions, red: solvent exposed) b) Structure and biophysical confirmation of Mac1-CACHE3-HO _1706_4 binding with SPR c) Structure and biophysical confirmation of Mac1-CACHE3-HO _1706_7 binding with SPR.

### Chemical series expansion in the REAL Space

Having identified a novel, solid ligand, we sought to identify active analogues to confirm the new chemical series of neutral Mac1 inhibitors by exploring different substituents interacting with the oxyanion hole. It is important to note that the x-ray structure of CACHE3-HI_1706_56 - Mac1 was not yet available at the hit expansion step of CACHE3, hence for this stage we relied solely on the computational pose of CACHE3-HI_1706_56. By performing a similarity search within the REAL Space database (44 billion molecules) followed by Glide SP docking and visual inspection, we identified 75 close analogues that were subsequently submitted for quotation to Enamine. Based on availability and CACHE3 budget restriction, we then prioritized 19 molecules for synthesis, 17 of which were successfully synthesized. 16 of the 17 analogues displayed dose-response inhibition of the binding of Mac1 to the ADP-ribose peptide (Supplementary Figure 1), and 12 were confirmed as NSP3 Mac1 binders via SPR (Table 1).

**Table 1.**
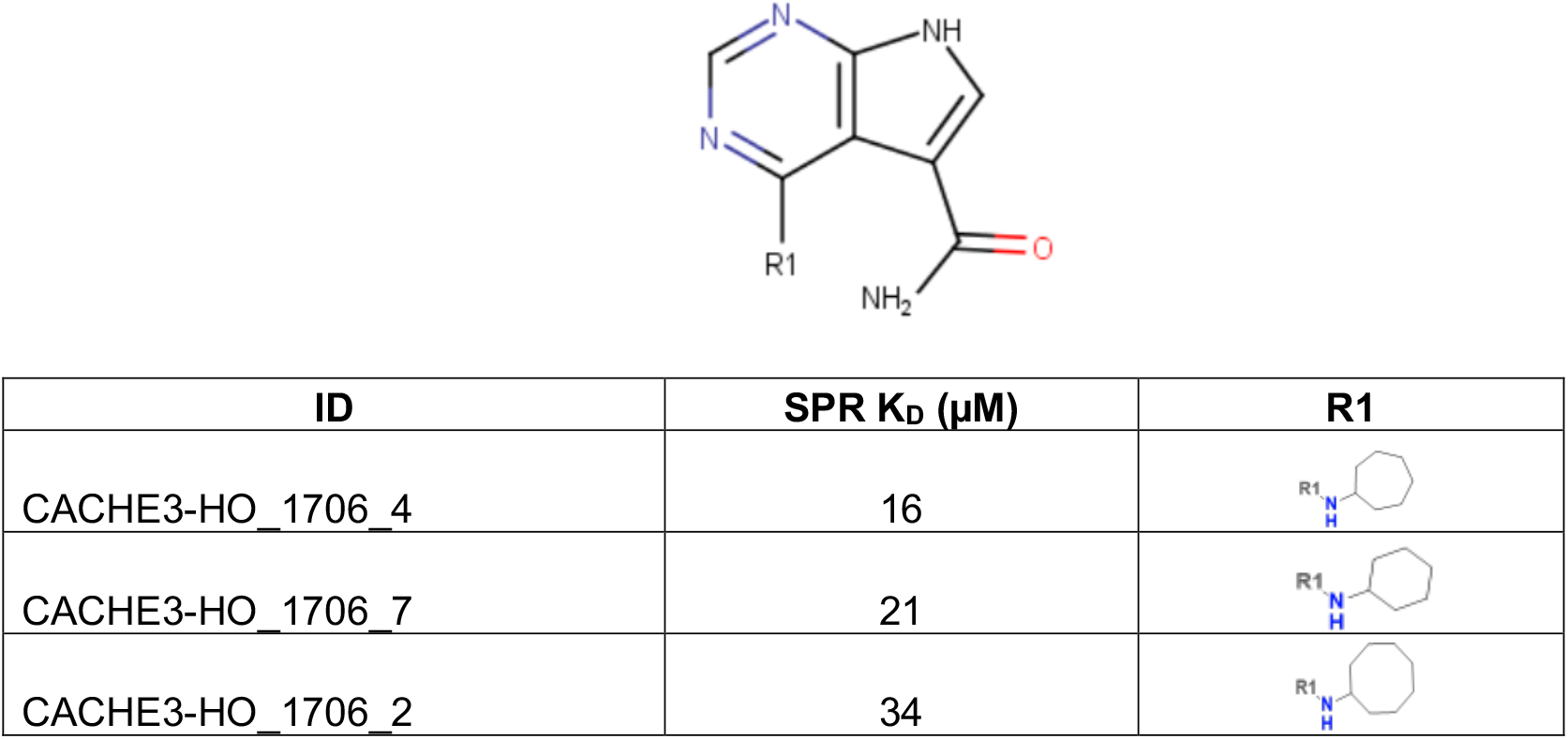
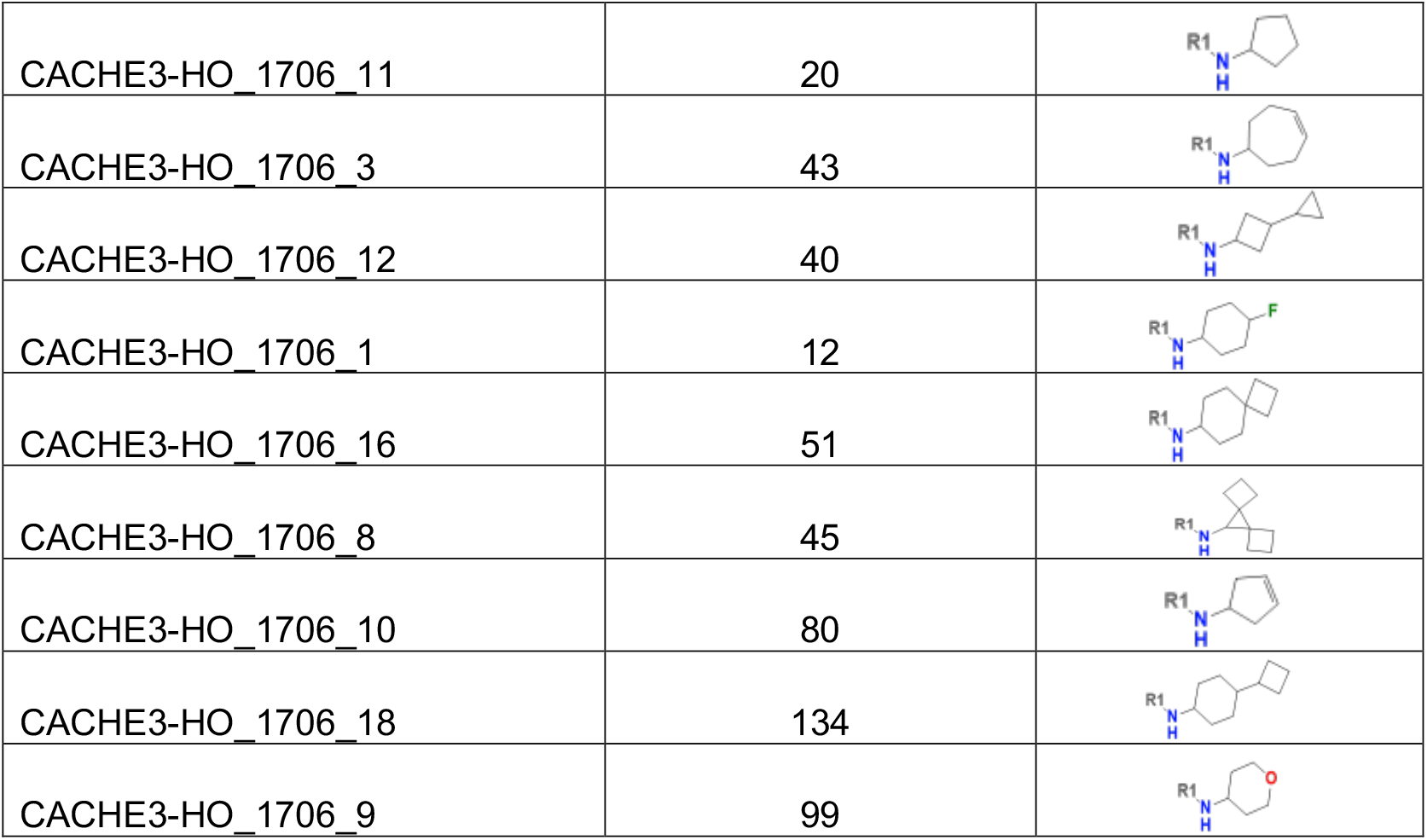
Structure-based exploration of CACHE3-HI_1706_56 by analogue search in Enamine REAL Space.

| ID | SPR K<sub>D</sub> (μM) | R1 |
| --- | --- | --- |
| CACHE3-HO_1706_4 | 16 |  |
| CACHE3-HO_1706_7 | 21 |  |
| CACHE3-HO_1706_2 | 34 |  |
| CACHE3-HO_1706_11 | 20 |  |
| CACHE3-HO_1706_3 | 43 |  |
| CACHE3-HO_1706_12 | 40 |  |
| CACHE3-HO_1706_1 | 12 |  |
| CACHE3-HO_1706_16 | 51 |  |
| CACHE3-HO_1706_8 | 45 |  |
| CACHE3-HO_1706_10 | 80 |  |
| CACHE3-HO_1706_18 | 134 |  |
| CACHE3-HO_1706_9 | 99 |  |

The selected analogues explored various neutral substitutions for the bromo-cyclohexene group including monocyclic alkane, mono cyclic alkene, spiro and dispiro alkanes to a small hydrophobic subpocket. Their predicted binding poses were consistent with the crystallographic structure of the original CACHE3-HI_1706_56 hit (Figure 4a). 5-, 6-, 7- and 8-membered monocyclic alkane were selected to probe the size effect of the rings on the activity, and 7-membered turned out to be the best group (K_D_=16 µM, Figure 4b, c and Supplementary Figure 1). Floro-hexane replacement demonstrated the overall best binding affinity with a K_D_ of 12 µM. Cyclopentene and cycloheptene, spiro[3.5]nonane and dispiro[3.0.35.14]nonane substitutions showed K_D_s in the range of 43 to 80 µM. The bridged compounds of 1,1’-bi(cyclobutane) and cyclobutylcyclohexane substitutions demonstrated lower activities corresponding to K_D_s of 51 and 134 µM, respectively. 9 molecules did not show any binding to PARP14a, confirming strong selectivity of the series toward NSP3 Mac1.

Importantly, our prioritization pipeline did not identify any CACHE3-HI_1706_56 analogue bearing a neutral hydrogen bond acceptor moiety placed in proximity of the oxyanion hole by docking, a modification that could be further explored to enhance target engagement.

## Conclusions

The expansion of tractable chemical space continues to challenge structure-based drug discovery due to the increasing computational costs. Active learning strategies have delivered consistent results in accelerating this exploration, yet it remains unlikely that such methods can be adapted efficiently to libraries exceeding billions of compounds. Fragment-based and combinatorial screening methods have also shown success, but their applicability across diverse targets, particularly those with challenging binding sites, remains relatively underexplored. A further obstacle is the amplification of artifact molecules due to the chemical congestion of typical of ultra-large spaces^38^.

In this study, we developed a bottom-up screening strategy based on AI-accelerated screening of a diverse, representative subset of molecules from an ultra-large chemical space to rapidly identify novel inhibitor chemotypes of SARS-CoV-2 NSP3 Mac1, which target engagement was unambiguously confirmed by x-ray crystallography. The identified hit was subsequently expanded through an ultra-large analogue search through 44-billion-compounds chemical space, yielding a novel chemical series with robust inhibitory and binding activity to Mac1. Thus, our target-agnostic approach effectively identified a novel series of neutral (noncharged) Mac1 inhibitors with potential for further optimization, and demonstrated competitive performance in the prospective CACHE3 challenge against a range of alternative screening methods, including fragment-based, generative, and deep learning workflows^39^.

Taken together, these results demonstrate that coupling virtual screening of smaller, diverse libraries with ultra-large analogue expansion offers an efficient way to interrogate extremely large, structurally congested chemical spaces.

## Supporting information

Supplementary x-ray statistics

## Data and Code Availability

Chemical structures and experimental data for the compounds tested in the CACHE3 challenge are available in Supplementary Table 1. The crystal structure of CACHE3-HI_1706_56 in complex with SARS-CoV-2 NSP3 Mac1 has been deposited in the PDB with ID 7HPW (https://doi.org/10.2210/pdb7HPW/pdb). The Deep Docking code is freely available at https://github.com/jamesgleave/DD_protocol.

## Acknowledgments

This work was supported by a Discovery Grant from the Natural Sciences and Engineering Research Council (NSERC) of Canada (RGPIN-2023-04129) and a startup grant from the University of Ottawa awarded to FG, a NSERC Discovery Grant (RGPIN-2024-04153) awarded to AC, and NIH GM145238 and U19AI171110 awarded to JSF. Experimental testing was supported by an Open Science Drug Discovery grant from Canada’s Strategic Innovation Fund (SIF Stream 5) administered by Conscience as well as by NIH grant 1U19AI171292-01 (READDI-AViDD Center), and was conducted at the Structural Genomics Consortium, a registered charity (no: 1097737) that receives funds from Bayer AG, Boehringer Ingelheim, Bristol Myers Squibb, Genentech, Genome Canada through Ontario Genomics Institute [OGI-196], Janssen, Merck KGaA (aka EMD in Canada and the US), Pfizer, and Takeda.

## Conflicts of Interest

FG is a co-founder and advisor of In Virtuo Laboratories. JSF is a consultant to and a shareholder of Vilya Therapeutics and Relay Therapeutics. These companies had no role in the design or conduct of the study; in the collection, analysis, or interpretation of the data; or in the preparation, review, or approval of the manuscript.

## Notes

### Summary of Updates

Fix typos and chemical structures of analogues.

