## Supplementary Material for "Structure-based discovery of inhibitors of Mac1 domain of nonstructural protein-3 of SARS-CoV-2 by machine learning-augmented screening of chemical space"

*^#^Equal contribution*

**Supplementary Figures**

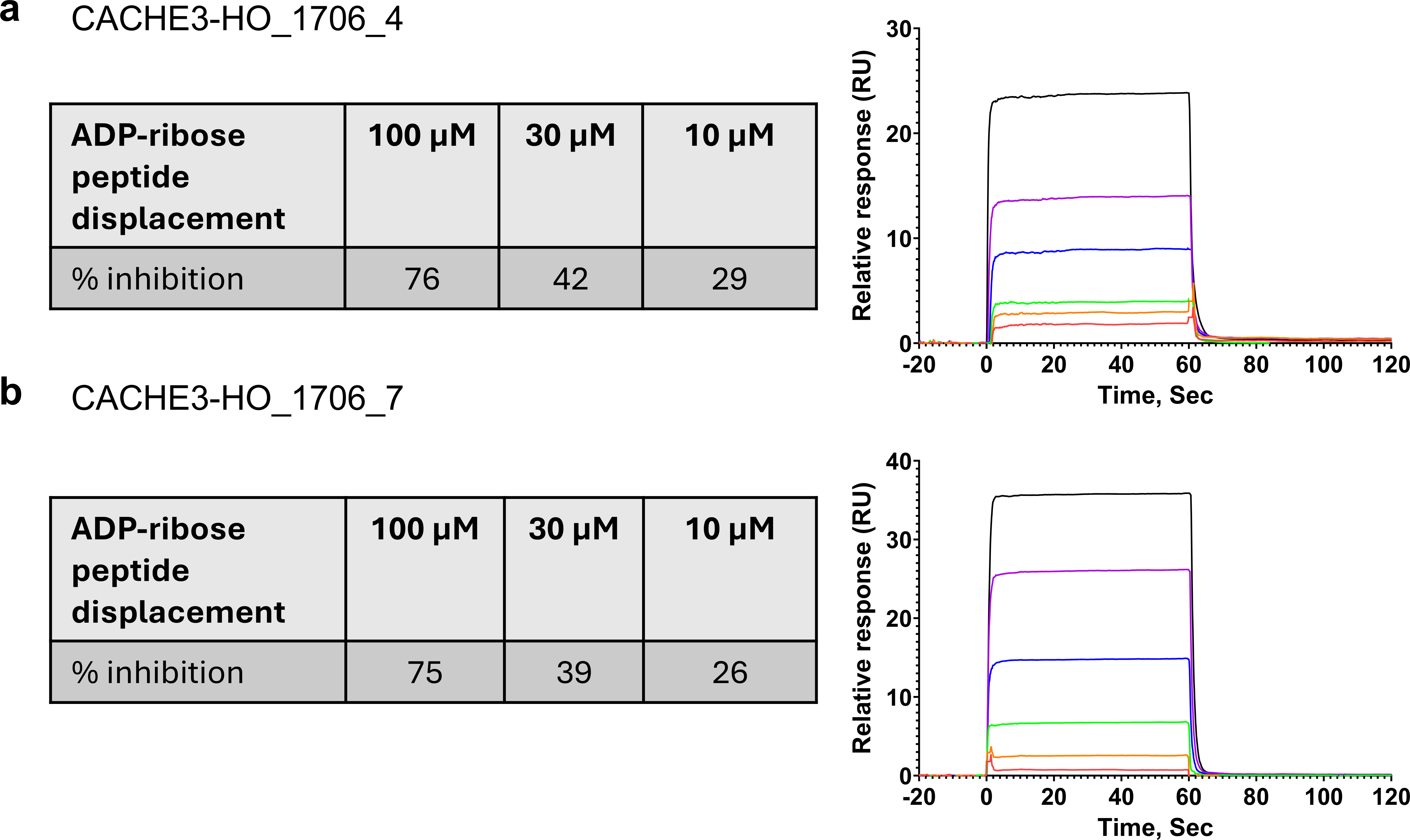

***Supplementary Figure 1.*** *ADP-ribose peptide displacement and SPR ligand binding confirmation for a) CACHE3-HO_1706_4 and c) CACHE3-HO _1706_7.*
